# Expanded transcriptomic analysis of human hepatic stellate cells links novel coding and noncoding products to human liver fibrosis

**DOI:** 10.1101/2022.02.01.478715

**Authors:** Amin Mahpour, Alan Mullen

## Abstract

End stage liver disease and liver failure occur primarily as a consequence of progressive fibrosis resulting from chronic liver injury. Hepatic stellate cells (HSCs) are the primary cell type responsible for production of the extracellular matrix (ECM) that forms the fibrotic scar. While the essential role of HSCs is understood, there remain no treatments that target HSCs to inhibit the development or progression of fibrosis. We have performed analysis of the transcriptome of human HSCs to define the long noncoding (lnc) RNAs expressed in this cell type, including many not previously annotated. Through analysis of full-length RNA transcripts, we identified additional lncRNAs that were not assembled by short reads. We also discovered new isoforms of proteincoding genes that encode amino acid sequences that are not present in annotated isoforms. Analysis of non-polyadenylated RNAs did not identify additional genes encoding long noncoding RNA transcripts, but did reveal the presence of hundreds of circular (circ) RNAs, including those with potential for translation. Incorporating these transcripts and genes into analysis of a published dataset of human liver fibrosis revealed the induction of lncRNAs, novel protein isoforms, and circRNAs associated with development of disease. These results identify RNAs and amino acid sequences expressed in HSCs and associated with human liver disease that may serve as therapeutic targets to inhibit fibrosis or biomarkers to benchmark progression of disease.

## Introduction

Hepatic stellate cell (HSC) myofibroblasts are the primary cell type responsible for the deposition of the extracellular matrix that forms the fibrotic scar in liver disease (Friedman et al. 1985; Maher and McGuire 1990; Mederacke et al. 2013). Expression of protein-coding genes has been mapped in HSCs in many studies to understand how these gene products contribute to fibrosis (De Minicis et al. 2007). Long noncoding (lnc) RNA expression has also been evaluated in limited capacities in human HSCs and has primarily relied on analysis of lncRNAs contained in reference annotations or evaluation of lncRNAs known to be expressed across many cell types (Gao et al. 2018; Zheng et al. 2015; Yu et al. 2015).

Analysis of RNA expression by RNA sequencing (RNA-seq) is primarily performed using short reads (25-150 nucleotides). Reads of 100-150 nucleotides can be used to assemble RNA transcripts by mapping reads to exons and splice junctions across exons. Previous studies that have assembled the transcriptome to identify the lncRNAs expressed in an individual cell type have primarily employed this approach (Cabili et al. 2011; Guttman et al. 2010). This approach can be powerful, but is also limited when there are regions of repetitive sequence or similar sequences shared in different parts of the genome or many different isoforms representing different combinations of exons (Weirather et al. 2017). Long read approaches that sequence the full RNA transcript are now being employed to improve accuracy of mapping. This approach generates full transcripts but is limited to fewer reads.

In addition to linear RNA products, genes can also produce circular RNAs (circRNAs)(Salzman et al. 2012; Jeck et al. 2013). These circRNAs have been shown to act as microRNA (miRNA) sponges, affect transcription, and encode protein products (Memczak et al. 2013; Y. Zhang et al. 2013; Wang and Wang 2015; Y. Yang et al. 2017). CircRNAs are produced from backsplicing in which the 3’-end of an downstream exon splices to the 5’-end of an upstream exon to generate a circle of single stranded RNA (Capel et al. 1993), and these products are most commonly produced from protein-coding genes (Salzman et al. 2012). Some circRNAs are found across multiple cell types while others appear to be more cell type specific (Salzman et al. 2013).

To better understand the RNA transcripts that are present in human HSCs, we applied RNA-seq analysis to short read polyadenylated and non-polyadenylated RNAs and full transcript analysis to polyadenylated reads. This analysis identified lncRNAs, circRNAs, and novel isoforms of protein-coding genes expressed in human HSCs. We then identified transcripts that were enriched in patients with liver fibrosis. Together, these analyses expand our understanding of the HSC transcriptome and reveal previously unannotated RNA products that could serve as therapeutic targets or biomarkers to support management of patients with chronic liver disease.

### Defining long noncoding RNAs in HSCs

We first performed RNA sequencing (RNA-seq) to define the linear, polyadenylated RNA transcriptome in human HSC myofibroblasts using 100 nucleotide paired-end (100PE) directional high-depth sequencing. Using a genome-guided approach, reads were assembled into a transcript model (Figure 1A). This transcript model identified 123,948 transcripts that map into 48,826 different loci. We detected transcripts arising from 12,857 annotated protein-coding genes, which included 66,199 protein-coding isoforms (Figure 1B). The analysis also identified 18,254 isoforms encoded by 9,146 protein-coding genes that were not previously annotated (Gencode v38).

**Figure 1.**
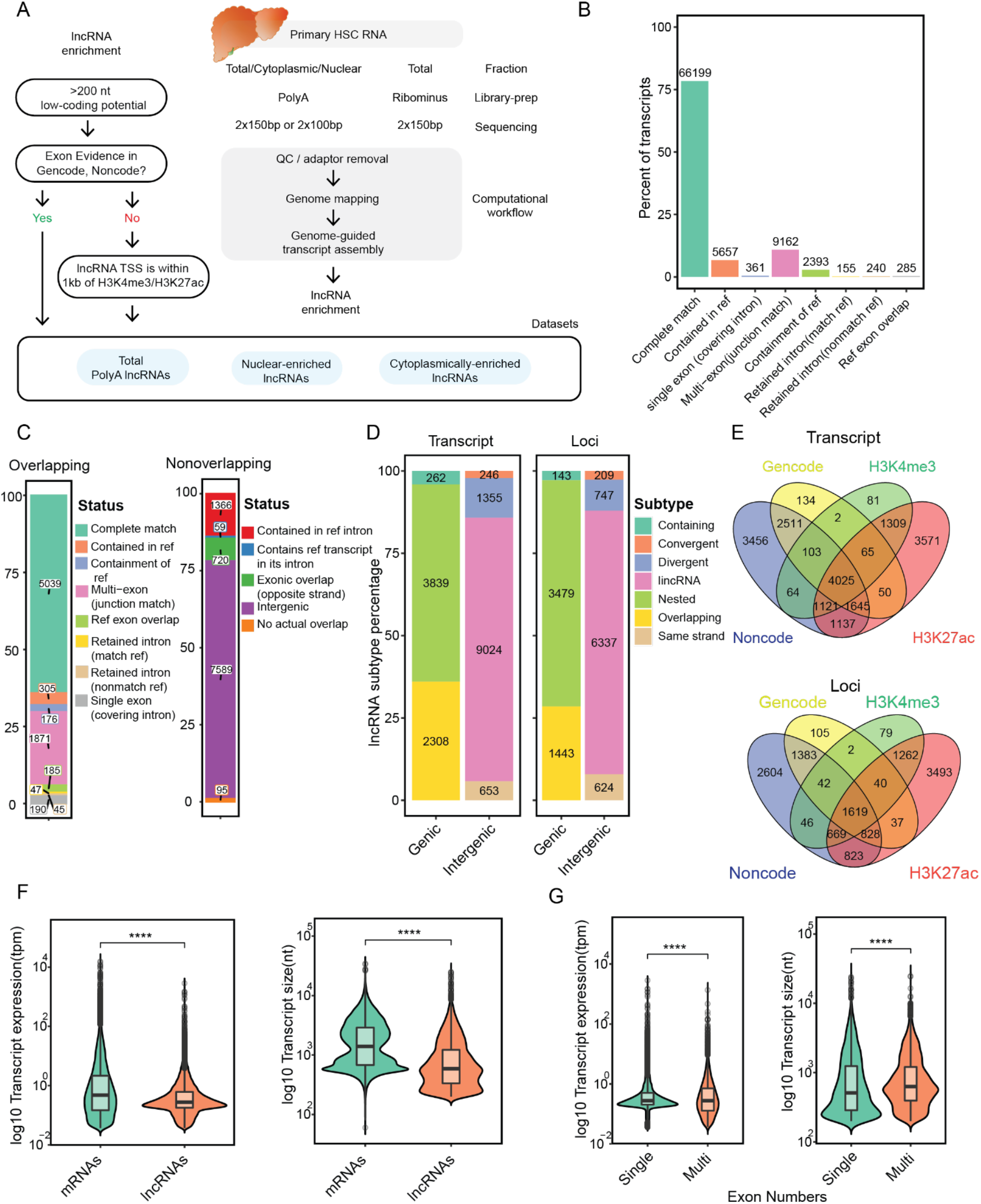
Defining lncRNA transcripts in primary human HSCs. **A)** Schematic of computational pipeline to generate HSC lncRNA datasets from RNA-seq libraries produced from HSC myofibroblasts. **B)** Assembled transcripts that either fully or partially overlap with proteincoding genes. More than 75 percent of assembled transcripts fully match with protein-coding transcripts annotated in Gencode. **C)** Comparison of assembled HSC lncRNA loci to those annotated in Gencode. The comparison is divided into HSC lncRNAs that overlap Gencode lncRNAs (left) and those that do not overlap with Gencode lncRNAs (right). The percentage of lncRNA loci in each classification is indicated on the y-axis and the number of loci is included on the plot. **D)** Classification of lncRNAs transcripts and loci in relation to protein-coding genes. The percentage of lncRNA transcripts (left) and loci (right) in each classification is indicated on the y-axis and the number of loci is included on the plot. Most assembled lncRNAs are not associated with a protein-coding transcript and are labeled as intergenic. **E)** Venn diagram showing how lncRNAs defined in HSCs are distributed across annotations in Gencode, Noncode and overlap at transcription start sites (TSS) with H3K4me3 and H3K27ac peaks. Analysis was performed for lncRNAs at the level of loci (top) and transcripts (bottom). The number of loci/transcripts is indicated for each region. **F)** Transcript expression (log10 of transcripts per million, tpm) is displayed for protein-coding genes andlncRNAs on the left. Transcript length (nucleotides, nt) for protein-coding and lncRNA genes is shown on the right. **G)** Transcript expression for single and multi-exon lncRNAs transcripts on the left, and transcript length is shown for single and multiexon lncRNAs on the right. Wilcoxon signed-rank test was used to determine statistical significance. **** p-value<0.0001.

We next defined the transcripts that represent lncRNAs. First, we filtered out transcripts that share one or more exons with annotated protein-coding genes. Next, we removed transcripts that are less than 200 nucleotides in length. We then filtered the transcripts to identify those with low protein-coding potential based on intrinsic features of transcript sequence (Kang et al. 2017). We compared our lncRNA dataset with lncRNAs annotated in Gencode (Derrien et al. 2012) (Figure 1C) and Noncode (Zhao et al. 2016) (Figure S1A). LncRNAs from HSCs that share one or more exons with annotated lncRNAs from Gencode or Noncode were included in the final list of lncRNAs.

Histone 3 lysine 4 trimethylation (H3K4me3) marks active promoter regions (Guttman et al. 2009; Mikkelsen et al. 2007) and histone 3 lysine 27 acetylation (H3K27ac) marks active enhancers (Creyghton et al. 2010). We performed Cut & Tag to determine sites of H3K4me3 and H3K27ac in HSCs. The remaining transcripts that passed all previous filters but were not contained in Gencode or Noncode were included as lncRNAs if their transcription start sites were associated with H3K4me3 and/or H3K27ac. Through this analysis, we defined 17,687 transcripts encoded by 12,434 loci that were distinct from protein-coding loci, demonstrated low protein-coding potential, and were greater than 200 nucleotides in length.

Of the total lncRNAs defined in HSCs (17,687 transcripts mapping to 12,434 loci), we identified 7858 lncRNA transcripts belonging to 3818 loci that are annotated in Gencode. Of the remaining lncRNAs, we identified 4868 of transcripts corresponding to 3822 loci that were in Noncode. This leaves 4,870 lncRNA loci that are uniquely defined in the HSC analysis (Figure 1E). In agreement with previous studies (Derrien et al. 2012), lncRNAs are expressed at lower levels than protein-coding genes, and tend to have shorter transcript lengths compared to protein-coding genes (Figure 1F). Within lncRNA transcripts, single exon lncRNAs are expressed at higher levels and have shorter lengths than multi-exon lncRNAs (Figure 1G).

Noncoding transcripts arising from H3K27ac-modified chromatin are often referred to as enhancer RNAs (eRNAs) (Melgar, Collins, and Sethupathy 2011). While many eRNAs are short, unspliced, non-polyadenylated, and bidirectionally transcribed, unidirectional eRNAs are polyadenylated and can be spliced (Koch et al. 2011). Of the lncRNAs identified in HSCs, 4,335 loci are classified as unidirectional eRNAs (Table S1).

In the process of defining lncRNAs, we filtered out transcripts that are predicted to have protein-coding potential. We identified 933 loci (1,556 transcripts) that met all other criteria for lncRNAs except they were predicted to have protein-coding potential and are not annotated in Gencode or Noncode (Table S1). Further investigation will be required to understand the functions of these gene products and whether or not they encode proteins.

LncRNAs are frequently annotated from polyadenylated RNA libraries (Guttman et al. 2010), but other noncoding RNAs over 200 nt in length have been described that are not poly-adenylated (L. Yang et al. 2011). We expanded our analysis to try to capture additional genes expressed in HSCs by assembling transcripts from ribosomal RNA (rRNA)-depleted RNAs to capture long noncoding transcripts that may not be polyadenylated. We analyzed 100PE directional reads using the same pipeline that we used for polyadenylated RNAs. We identified 1,272 lncRNA transcripts representing 1,132 loci, and all of these loci were also detected in the polyA data (Table S1). The rRNA-depleted libraries contain both polyadenylated and non-polyadenylated (nonPolyA) RNAs. We cannot assess the fraction of nonPolyA transcripts with this analysis, but we did not find evidence of additional genes expressing nonPolyA transcripts that met all other criteria for lncRNAs.

### Long-read sequencing reveals the transcriptome diversity in HSCs transcripts

RNA transcript assembly is commonly performed using short reads of ~100 nucleotides to annotate mRNAs and lncRNAs, but transcripts assembled from these reads can fail to capture the true isoform diversity of a given transcriptome and could lead to inaccurate predictions of protein-coding transcript isoforms expressed in HSCs (Tørresen et al. 2019). The long-read sequencing platform is increasingly gaining popularity to study complex biological samples such as cancers (Veiga et al. 2022). To overcome these deficiencies, we sequenced full-length polyA transcripts in HSCs. We employed Single-Molecule Real-Time sequencing (SMRT-seq) followed by the Pacbio Iso-seq computational pipeline to generate full-length PolyA transcripts (Figure 2A).

**Figure 2.**
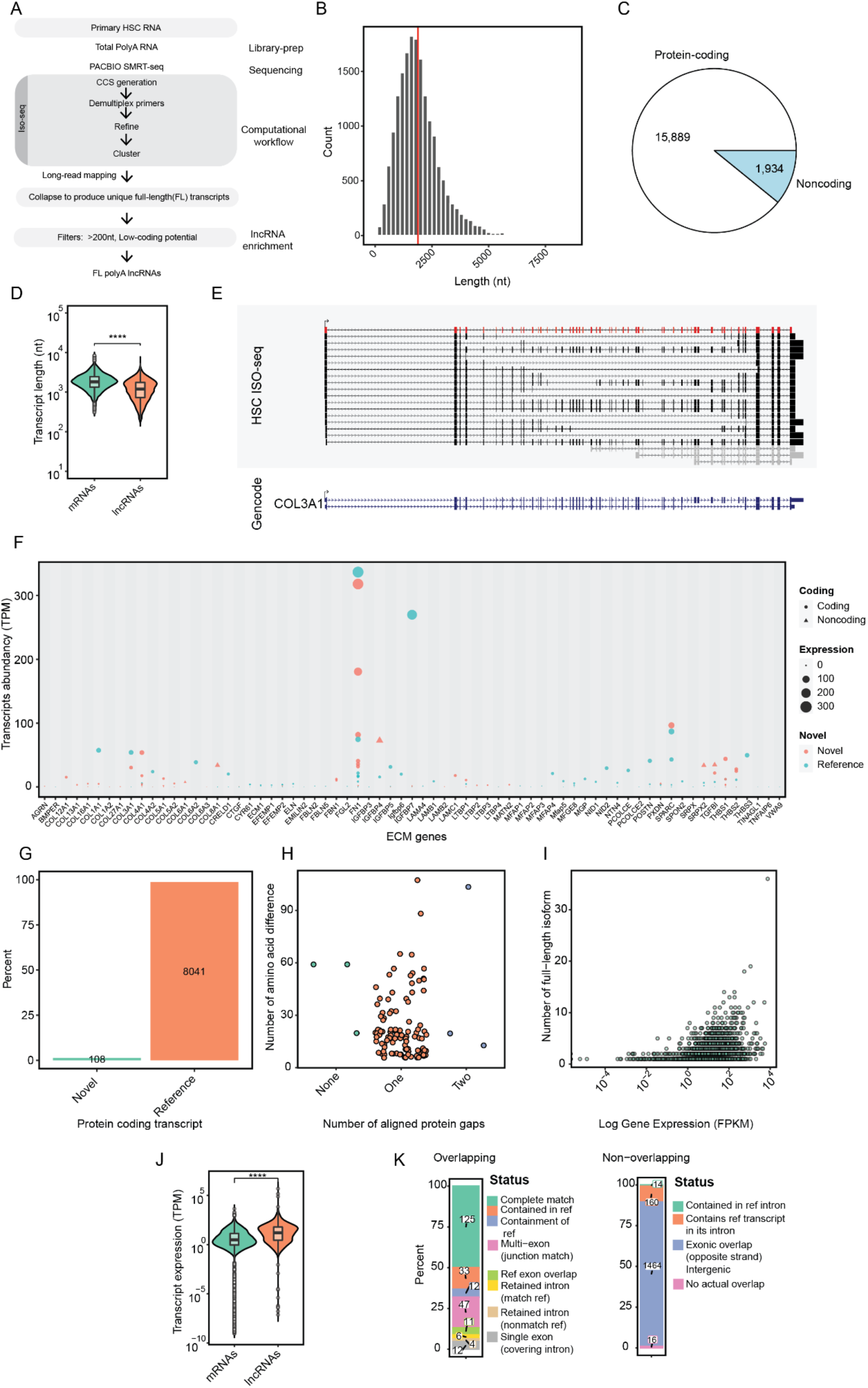
Annotating protein-coding and lncRNA transcripts with long read sequencing. **A)** Schematic showing individual steps in processing of full-length transcripts from SMRT-seq sequencing data. **B)** Size distribution of full-length HSC transcripts produced by the Iso-seq pipeline. The vertical red line indicates the mean length size of transcripts. **C)** Venn diagram indicates the distribution of full-length protein-coding and lncRNA transcripts. **D)** Transcript length (nt) is indicated on the y-axis for coding and lncRNA transcripts. **E)** Isoforms identified for *COL3A1* are shown. The transcript in red indicates the long-read transcript that matches the bottom reference transcript annotated in Gencode (bottom). Transcripts marked in gray are shorter transcripts that do not share 5’-ends with reference transcripts and are excluded from downstream analysis. **F)** Expression of transcript isoforms belonging to extracellular matrix (ECM) genes. Green circles indicate annotated isoforms and orange circles indicate novel isoforms. The size of the circle indicates the level of expression, which is also quantified on the y-axis. Triangles indicate noncoding transcripts. **G)** Each novel isoform was evaluated to identify those that generate at least five contiguous amino acids that are different from all annotated isoforms. These isoforms are defined as novel and are plotted in comparison to the remaining protein-coding transcripts annotated by long-read sequencing. **H)** The number of novel amino acids (y-axis) are plotted versus the gaps in alignment (x-axis) for each of the 108 isoforms identified in (G). Single gaps are represented in orange while other no gaps (extension of sequences at the 3’-end) in green and double gaps are represented in blue. **I)** The number of transcript isoforms (y-axis) are plotted versus gene expression (Fragments Per Kilobase of transcript per Million mapped reads, FPKM) (x-axis). FPKM was calculated for each gene based on short-read data. **J)** Transcript levels (TPM, y-axis) are plotted for mRNAs and lncRNAs identified by long reads. Expression represents the number of full-length reads assembled for each transcript. This observation might be due to the sequencing bias for the transcripts that are highly expressed. Wilcoxon signed-rank test was used to determine statistical significance. **** p-value <0.0001. **K)** Comparison of lncRNAs annotated by full length transcripts in HSCs with those annotated by short-reads. The comparison is divided into lncRNA transcripts assembled by long-read sequencing in HSCs that overlap with those annotated by short-read sequencing and those that do not overlap with lncRNAs assembled by short read transcripts. The percentage of lncRNA transcripts in each classification is indicated on the y-axis and the number of transcripts is included on the plot.

Libraries were prepared to capture transcripts less than 5,000 nucleotides (nt) in length. We find that 90% of transcripts are between 500 and 5,000 in length with a mean of 1,916 nt, with the largest transcript containing 8507 nt (Figure 2B). Of the 17,823 transcripts identified, 15,889 are protein-coding and 1,934 are lncRNAs (Figure 2C), and we again observed that lncRNAs tend to be shorter in length than protein-coding transcripts (Figure 2D).

Analysis of individual genes reveals examples where many isoforms of a gene are detected that are not annotated in Gencode. While almost 64 percent of protein-coding of transcripts detected are annotated in Gencode, 36 percent contain isoforms that are not currently annotated. As an example, *COL3A1* has two annotated isoforms, while one annotated isoform plus 16 additional isoforms are detected in the long read data (Figure 2E). The variations in isoforms observed in *COL3A1* occur from skipped exons and novel splice junctions. An even more dramatic example is observed with the gene encoding COL1A1 (Figure S2A). We also confirmed the presence of unannotated splice junctions in *COL1A1* in HSCs using reverse transcriptase (RT) followed by amplification by PCR (Figure S2B). These results suggest that many unrecognized spice variants for genes are expressed, which can be challenging to annotate from short read data.

HCSs are the primary cell type responsible for production for the extracellular matrix (ECM), which forms the fibrotic scar in liver disease, and we next evaluated genes associated with this pathway. We identified 227 transcripts for 69 genes that are associated with ECM. Some genes such as *FN1*, *COL4A1*, *SPARC*, and *TGFBI* express novel isoforms with comparable expression to annotated isoforms (Figure 2F), while many other genes express novel isoforms that are less abundant than annotated isoforms. These findings suggest that there may be a greater diversity in protein isoforms composing the fibrotic scar than previously appreciated.

We next analyzed the novel splice variants to identify those that represent changes in amino acid sequence. Through analysis of protein alignment, we identified isoforms with novel amino acid sequences for 108 transcripts that correspond to 92 protein-coding genes (Figure 2G). These novel amino acids can introduce gaps or extend the sequence of reference protein products, most of which involve a single gap in alignment (Figure 2H). For this analysis, we required at least five contiguous new amino acids compared to annotated isoforms. The number of isoforms detected increases with gene expression (Figure 2I), which suggests that we are not yet at saturation with our current analysis, and additional isoforms will be identified with additional depth of sequencing.

We next evaluated lncRNAs identified from the full-length transcript data. Over 125 lncRNAs transcripts were identified that matched 98 loci that we already defined with the short read annotations in HSCs. More than 1,464 lncRNA transcripts mapping to 1,094 loci were also identified that were not annotated by short read analysis. We also compared the expression of full-length transcript by using short read RNA-seq data from HSCs. We find that lncRNAs detected in full-length long read sequencing are expressed in higher levels than the protein-coding genes, which might be the result of long read sequencing bias for highly expressed lncRNAs (Figure 2J). Compared to the previously annotated lncRNAs in HSCs, most of the transcripts defined from long read were intergenic and they did not overlap with annotated protein-coding or lncRNA loci (Figure 2K). These results indicate that assembly of short reads can identify many genes, but additional transcripts are still overlooked.

### Sequences involved with nuclear retention of lncRNAs

LncRNAs can be nuclear or cytoplasmic, while mRNA transcripts are predominantly cytoplasmic (Guttman et al. 2013). We next performed RNA sequencing on polyadenylated RNAs from nuclear and cytoplasmic fractions in HSC myofibroblasts to define the nuclear and cytoplasmic distribution of lncRNAs at the level of gene products and individual isoforms.

We first evaluated the efficiency of nuclear and cytoplasmic fractionation by assessing enrichment of mitochondrial RNA in the cytoplasmic fraction and nuclear lncRNAs, including *MALAT1*, *NEAT1* and *MEG3* in the nuclear fraction. The distinct separation of mitochondrial RNA to the cytoplasmic pool and *MALAT1*, *NEAT1* and *MEG3* to the nuclear pool indicates effective separation of nuclear and cytoplasmic fractions (Figure S3A).

We next evaluated the distribution of RNAs expressed from protein-coding and lncRNA genes. We find that a similar number of protein-coding genes are enriched in cytoplasmic and nuclear fractions (2179 and 2270 transcripts respectively) (Figure 3A). We next evaluated the distribution of lncRNAs at loci (Figure 3B) and transcripts levels (Figure 3C). Through this analysis, we find that a larger fraction of lncRNA loci (1,618 nuclear loci and 1,250 cytoplasmic-enriched lncRNA loci) and transcripts (2,184 nuclear vs 1,693 cytoplasmic-enriched lncRNA transcripts) are nuclear-enriched. The significant majority of transcripts expressed from the same loci were enriched together in the nucleus or cytoplasm (Figure 3D). LncRNAs that predominantly localize to the cytoplasm tend to be shorter in total length (Figure 3E), while they are expressed at a higher level (Figure 3F) and contain more exons (Figure 3F) than lncRNAs that predominantly localized to the nucleus. These data suggest that while lncRNA transcripts tend to be enriched more in nuclear fraction, they are abundant in both the nucleus and cytoplasm.

**Figure 3.**
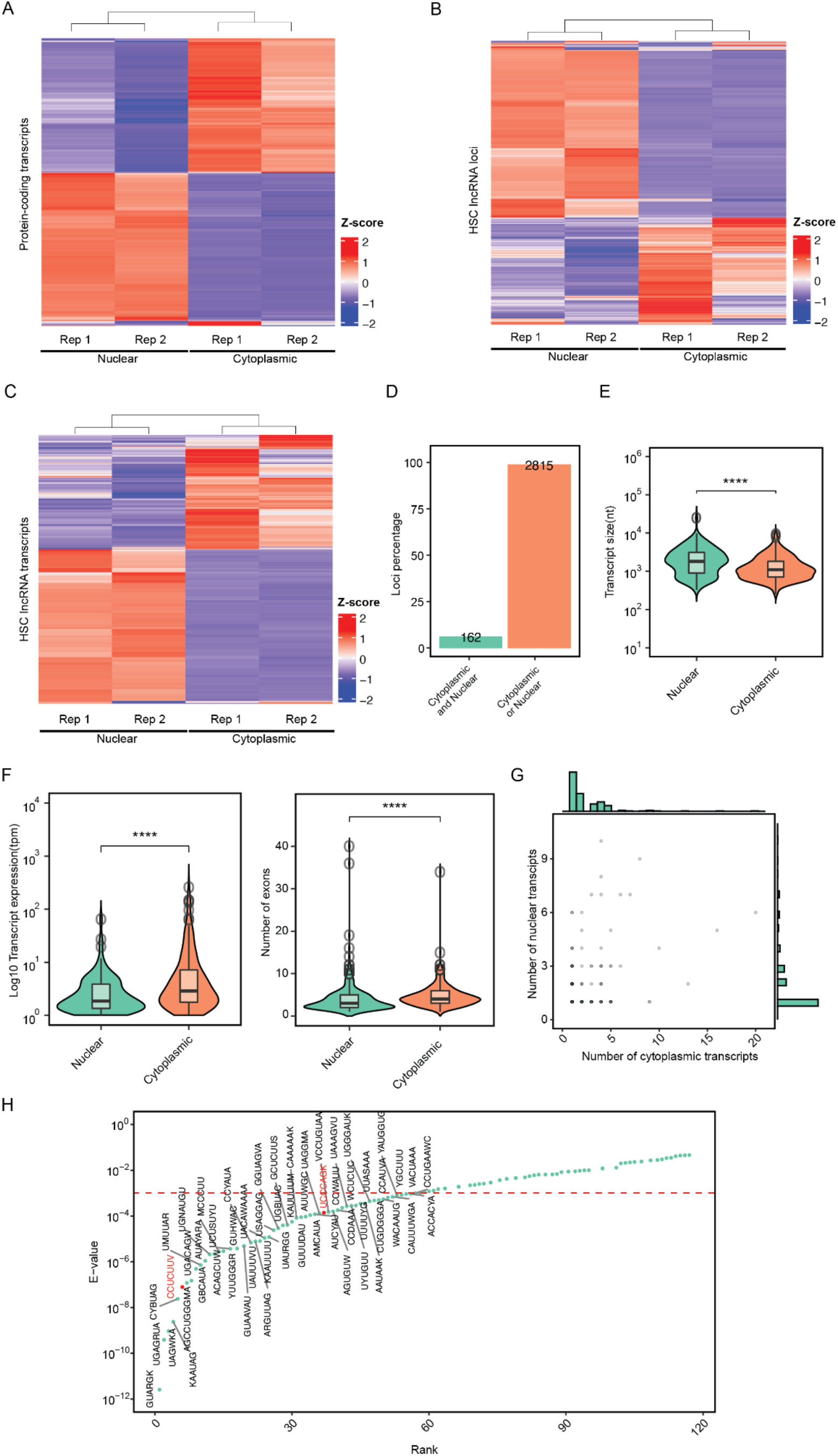
Nuclear and cytoplasmic localization of lncRNAs. **A)** Heatmap plot shows distribution of protein-coding transcripts across nuclear and cytoplasmic fractions in HSCs. To be considered enriched in a fraction, transcripts should meet the statistical criteria of adjusted p-value <0.05 and |Log2 Fold Change| >1. Row-scaling has been applied to the heatmap to show enrichment of transcripts/loci across fractions and replicates. **B)** Heatmap plot shows the distribution of polyA HSC lncRNA loci. **C)** Heatmap plot shows the distribution of polyA HSC lncRNA at the transcript level in C. **D)** The percentage of lncRNA loci (y-axis) showing transcripts distributed across both nuclear and cytoplasmic fractions (green, Cytoplasm and Nucleus) and those that are enriched in either nuclear or cytoplasmic fractions (orange, Cytoplasm or Nucleus). Loci are considered to produce transcripts that localize to both the cytoplasm and nucleus if at least one transcript from each loci meets statistical criteria (log10 adjusted p-value <0.05 and log2 fold change of 1) for enrichment in the nucleus and another transcript reaches criteria for enrichment in the cytoplasm. Loci are considered to produce transcripts that localize to the nucleus or cytoplasm if at least one transcript from each loci meets statistical criteria (p-value < 0.05 and log2 fold change of 1) for enrichment in either the nucleus or cytoplasm and no transcripts meet statistical criteria for enrichment in the other fraction. **E)** Transcript size (nt, y-axis) is shown for nuclear-enriched transcripts and cytoplasmic-enriched transcripts. Wilcoxon signed-rank test was used to determine statistical significance. **** p-value <0.0001. **F)** Expression (left) and number of exons (right) are plotted for nuclear and cytoplasmic-enriched lncRNA transcripts originating from the same loci. Wilcoxon signed-rank test was used to determine statistical significance. **** p-value <0.0001. **G)** The number of nuclear transcripts (y-axis) were plotted versus the number of cytoplasmic transcripts (x-axis) for lncRNA from the same loci. **H)** Differential enrichment analysis was performed to define motifs enriched in transcripts localized to the nucleus compared to transcripts localized to the cytoplasm and originating from the same lncRNA loci. Motifs in red are referenced in the text. Wilcoxon signed-rank test was used to determine statistical significance (p-value <0.0001) and is marked by a dotted red line.

A small fraction of lncRNA loci contain transcripts with differential localization between the nucleus and cytoplasm. RNA sequences have been reported that promote nuclear retention of lncRNAs (Ross et al. 2021), and we focused on these loci to evaluate the sequences that might determine localization. The most common example occurs with lncRNA loci that express two isoforms, and these examples are more weighted to lncRNA loci encoding more transcripts that localize to the cytoplasm than the nucleus (Figure 3G). We identified the sequence signatures that are specifically enriched in transcripts localized to the nucleus compared to transcripts from the same lncRNAs locus that were localized to the cytoplasm (Figure 3H). The most enriched motif was QUARGK for which we could not identify a binding protein in available databases. CCUCUUV is recognized by MATR3 (Ramesh et al. 2020), an RNA binding protein involved in paraspeckle assembly that regulates *NEAT1* expression (Banerjee et al. 2017). We also found enrichment of the HNRNPK binding motif (UCCCAGK) (Feng et al. 2019). This sequence resembles the SIRLOIN motif that was found to be enriched in Alu elements, is recognized by HNRNPK, and promotes nuclear localization (Lubelsky and Ulitsky 2018). These results show that lncRNAs retained in the nucleus of HSCs tend to be longer, and sequences enriched in these lncRNA transcripts may serve as binding sites for nuclear proteins.

### Widespread expression of circRNAs in HSCs

Recent studies have identified the presence of non-linear RNA products that can regulate cellular function. CircRNAs are formed through a backspling event where the 3’-end of an downstream exon is spliced to the 5’-end of an upstream exon to form a circular RNA product. To provide a deeper understanding of the potential role of circRNAs in HSC biology, we evaluated RNA-seq data from rRNA-depleted RNA to identify back splice events that define the population of circRNAs in HSCs (Figure 4A). We identified 8,284 unique back splice events that may produce circRNAs (Table S1). We filtered the list to 1,094 high-confidence circRNAs, identified by the presence of more than two different reads supporting the backsplice (Zhou et al. 2017). 1,048 circRNAs were produced from protein-coding exons, while 16 were produced from lncRNA exons. There is equal distribution of circRNAs from the Watson and Crick (plus and minus) strands (Figure 4B) and a mean of six reads support the circRNA transcripts identified (Figure 4B). The median number of exons in HSC circRNA is three and ranges from one to eighteen exons (Figure 4C). Analysis of the back splice junctions (BSJs) of HSC circRNAs reveals enrichment towards the 5’-end of genes (Figure 4D). We also find that the number of exons in a circRNA is poorly correlated to the number of exons in the parent gene (Figure 4E). To confirm the presence of identified circRNAs annotated by this analysis, we performed PCR using divergent primers that will only amplify the BSJ that create *circATXN1* and *circFAT1* (Figure 4F).

**Figure 4.**
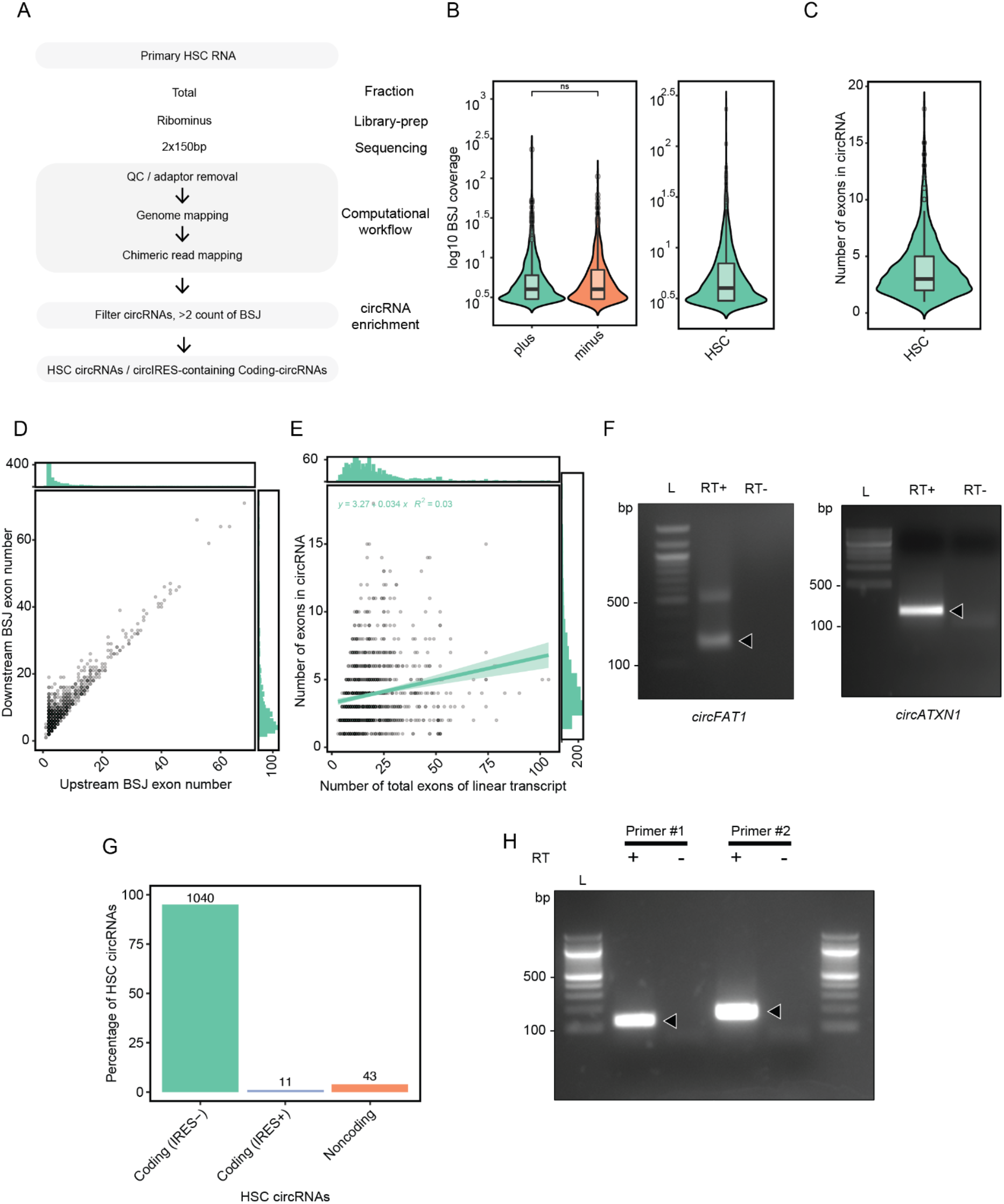
Defining circRNA in HSCs. **A)** Schematic showing steps toward generation of circRNAs from HSC cells. **B)** Distribution of circRNA as evaluated by backsplice junction (BSJ) quantification across plus (Watson) and minus (Crick) strands of the genome. **C)** The number of exons that are contained in HSC circRNAs is plotted and varies from 1 to 18. **D)** Plot indicating the exon numbers for upstream and downstream exons that contribute to the formation of BSJ. **E)** The number of exons contained in each circRNA (y-axis) are plotted versus the number of exons in the linear transcript of the parent gene (x-axis). The number of exons in the parent gene was defined as the number of exons in the isoform with the most exons. **F)** RT-PCR was performed on total RNA from HSCs. Divergent primers that only amplify across the BSJ of *circFAT1* (left) and *circATXN1* (right) were used for amplification. DNA amplicons were visualized on agarose gel. White arrows indicate the predicted size of the amplicon. Ladders are located on the left of each gel, and the most relevant size marked on the ladder (bp) is indicated to the left of the gel. RT+ indicates total RNA that underwent reverse transcription and RT-indicates total RNA that did not undergo reverse transcription prior to PCR. **G)** CircRNAs were assessed to identify those with open reading frames (coding). CircRNAs with open reading frames were then divided into those that contain a circIRES (Coding, IRES+) and those that do not contain a circIRES (Coding, IRES-). The number of circRNAs that originate from lncRNAs was also plotted for comparison. The percentage of circRNAs falling into each group out of the total circRNAs defined is shown on the y-axis. The total number of circRNAs in each category is indicated on each bar of the plot. **H)** Circular form of ELK4 (*circELK4*) is among the only two circRNA whose coding sequence is fully preserved and RT-PCR was performed to confirm the presence of this lncRNA in HSCs. Two sets of primers (Primers #1 and Primers #2) were used to amplify across the predicted BSJ. Arrows indicate predicted amplicon size. RT-indicates total RNA that did not undergo reverse transcription before amplification. Ladders are present in the first and last lanes of the gel.

We next evaluated circRNAs for protein-coding potential by scanning for sequences that could act as internal ribosomal entry sites (Chen et al. 2021). We identified 1,051 circRNAs with a sequence containing an open reading frame suggestive of possible translation (refer to methods), and two of these circRNAs contain a full open reading frame (ORF) for the parent gene (Table S1). To test the existence of example circRNAs that might have protein-coding potential, we amplified across the BSJ predicted for *ELK4*, which is predicted to form a circRNA containing the full *ELK4* coding region (Figure 4H), and by amplifying with two sets of divergent primers, we cloned and confirmed the existence of the full circular RNA by Sanger sequencing (Figure S4A). CircRNA-specific internal ribosomal entry sites (circIRES) are RNA sequences that allow ribosomal recruitment to circRNA and translation of open reading frames within the circRNAs to produce protein products (Chen et al. 2021). We next evaluated these 1051 circRNAs to identify those containing circIRES sequences identified to promote translation in circRNAs upstream of ORFs (Chen et al. 2021). We identified 11 circRNAs expressed in HSCs with high protein-coding potential based on the presence of an ORF and a circIRES predicted to promote translation in circRNAs. Taken together, these data show that circRNAs are abundantly expressed in HSCs, and circRNA products include those that have protein-coding potential.

### Expression analysis of HSC transcripts in liver fibrosis patients

HSC myofibroblasts are the primary cell type responsible for the production of the ECM that leads to fibrosis in chronic liver disease. We next applied our deeper understanding of the HSC transcriptome to evaluate changes that occur with the development of liver fibrosis in patients with non-alcoholic fatty liver disease (NAFLD) (Pantano et al. 2021).

We first evaluated lncRNA expression in total liver across a range of fibrosis levels associated with NAFLD (Figure 5A). RNA-seq data was analyzed from liver samples of individuals with no steatosis, those with steatosis and no fibrosis (F0), and those with steatosis and different stages of fibrosis (F1-F4), with F4 representing cirrhosis (Pantano et al. 2021). We identified 251 lncRNA transcripts that are enriched with more severe liver fibrosis, and 92 of these lncRNAs represent transcripts that were previously not annotated in Gencode. These results show that there are individual lncRNAs that are expressed by HSC myofibroblasts and enriched with the development of liver fibrosis associated with NAFLD.

**Figure 5.**
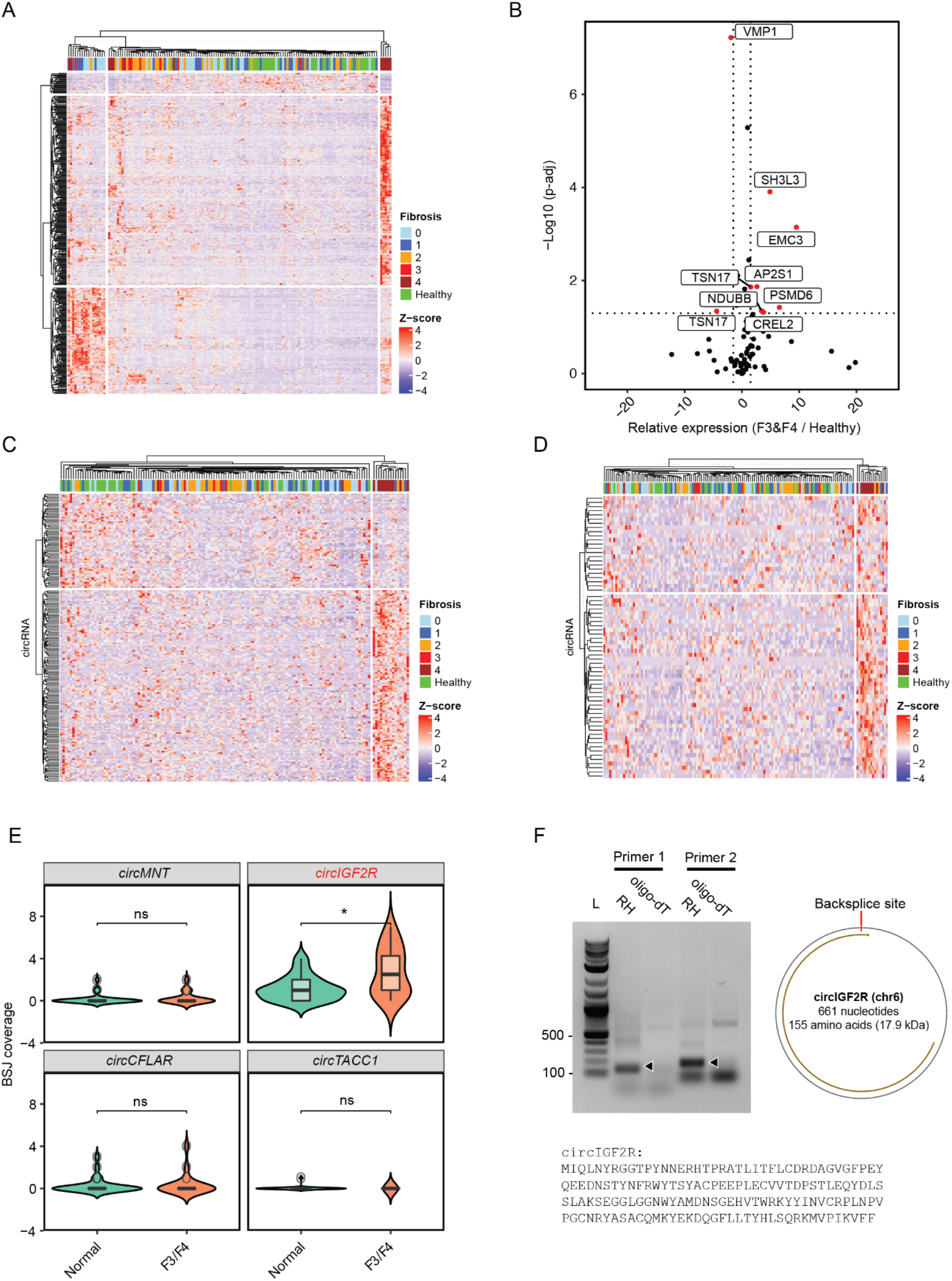
Expression of RNA products with NALFD fibrosis. **A)** Expression analysis of HSC lncRNAs in patients diagnosed with NAFLD with different degrees of fibrosis (F0-F1). Non-hierarchical clustering was performed to identify lncRNAs present in HSCs that are enriched with development of fibrosis. Z-scores indicate enrichment of expression, and the stage of fibrosis (F0-F1 are indicated across the top). Green indicates samples with normal histology. **B)** Relative expression of transcripts with altered amino acid sequence to that of transcripts that code for reference protein in normal samples versus F3/F4 samples. The y-axis indicates the log10 p-adj value and the x-axis indicates fold enrichment. Red dots indicate isoforms that are enriched by both adjusted p-value (0.05) and fold change (2) thresholds. **C)** Differential expression analysis of BSJ reads in patient samples for all the circRNA defined in data from human samples that show differential expression. **D)** Differential expression of circRNAs defined in HSCs across human samples. **E)** Expression analysis of BSJ belonging to four HSC circRNAs that contain both coding sequence and circIRES and are detected in the NAFLDpatient dataset. For each circRNA the number of BSJs detected (y-axis) is plotted for patients with normal histology and F3/F4 fibrosis. Only *circIGF2R* shows a statistical enrichment with the development of fibrosis. Wilcoxon signed-rank test was used to determine statistical significance. * p-value<0.05. **F)** PCR validation of *circIGF2R* using two different primer sets that will only amplify across a BSJ. Arrows on the agarose gel indicate expected product size. We used two different primers, RH (random hexamers) and oligo-dT to generate template cDNAs for this analysis. Oligo-dT primes linear polyA-containing transcripts and hence circRNAs BSJ are not expected to be amplified. The ladder (L) is located in the first lane, and the 500 bp and 100 bp markers are indicated on the left for reference. A model of the predicted product is indicated on the right (with additional details in Figure S4B). The predicted amino acid sequence encoded by *circIGF2R* is shown at the bottom.

We identified 108 protein isoforms that had not been described previously that produce amino acid sequences that are unique from those expressed from annotated isoforms of the same gene (Figure 2G). We next used the RNA-seq data in human samples to ask if any of these isoforms are induced in F3-F4 fibrosis compared to healthy liver samples (Figure 5B). Of the 108 isoforms, we identified seven produced from seven different genes that are enriched with the development of fibrosis. These results suggest that there are previously unannotated isoforms of proteins with unique amino acid sequences that are induced with development of fibrosis in NAFLD.

We then evaluated expression of circRNAs identified in HSC myofibroblasts with the development of liver fibrosis. We were able to perform this analysis because the RNA-seq analysis for NAFLD patients was performed on rRNA-depleted rather than polyA-enriched RNA (Pantano et al. 2021). We first identified all BSJ in patient samples, and then identified BSJs that are enriched in F4 fibrosis (Figure 5C). We next repeated the analysis with HSC circRNAs and produced a list of 56 circRNAs that are enriched with F4 fibrosis (Figure 5D). Finally, we analyzed expression of the circRNAs containing circIRES sequences shown to promote translation in HSCs. Only four were detected in the human data and one circRNA, encoded by *IGF2R* was enriched in patients with F3-F4 fibrosis compared to those with normal liver histology (Figure 5E). RT-PCR was performed using two pairs of divergent primers that only amplify the circRNA to confirm the presence of circIGF2R in HSC myofibroblasts. These results suggest that circRNA products are enriched with liver fibrosis in NAFLD, including those that have the potential to encode protein products.

## Discussion

Liver fibrosis is caused by chronic injury and activation of HSC myofibroblasts, the primary cell type responsible for production of the ECM proteins that form the fibrotic scar. While studies of gene expression in HSCs can now take advantage of broad annotations of protein-coding and lncRNA genes included in Gencode and additional lncRNA annotations available through platforms such as Noncode, these databases do not include the full range of RNA products produced by HSCs that could regulate their activity in fibrosis. To expand our understanding of gene products in HSCs, we analyzed polyadenylated and non-polyadenylated RNA products greater than 200 nt in length and performed additional analysis of full-length polyadenylated RNAs. This analysis in primary human HSCs allowed us to define novel transcripts associated with lncRNAs, novel isoforms of mRNAs, and circRNAs. We further linked these gene products to their expression across different stages of fibrosis in NAFLD to identify those that are more likely to mark and/or promote liver fibrosis.

This analysis identified 4,870 lncRNAs expressed in HSCs that were not previously annotated in addition to identifying expression of 3,818 lncRNAs contained in Gencode and 7,426 lncRNAs contained in Noncode. Analysis of NAFLD samples identified 251 lncRNAs that increased with fibrosis. Applying Gencode alone would have identified 124 of these and together with Noncode 182, but many of the lncRNA transcripts identified to increase with fibrosis came from those expressed in HSCs that were not previously annotated in other databases.

We identified 108 protein isoforms associated with presence of at least five contiguous amino acids that are not present in any other isoforms of the same proteins. This length was selected as the minimal number of amino acids required to form a full epitope to be recognized by an antibody (Buus et al. 2012), but changes in two to four amino acids could also affect protein function and could create unique sequence targets for antisense oligonucleotide depletion or novel protein epitopes that could be recognized by new antibodies. Single nucleotide polymorphisms (SNPs) would not affect our current analysis, and studies of additional donor HSCs will be helpful to establish the diversity and universality of the current findings.

CircRNAs are increasingly recognized for their biological activity, and recent work suggests that in addition to functioning as RNAs, circRNAs can also be translated into protein products (Chen et al. 2021). We identified 1,094 circRNAs expressed in human HSCs and found that 10 were increased at least by two fold in whole liver samples from patients with fibrosis in the setting of NAFLD (Table S1). The circRNA encoded by *IGF2R* was identified as the most likely candidate circRNA to be translated that was induced with development of severe fibrosis (Figure S4B). The *circIGF2R* product does not contain the full length *IGF2R* transcript, and the epitopes for antibodies that recognize the IGF2R protein are not contained in the circRNA product, limiting the ability to confirm expression of this isoform in human liver samples. *circIGF2R* instead encodes a fragment containing the Mannose-6-phosphate receptor homology domain, which could act as a dominant negative IGF2R, similar to the phenotype described for *circFGFR1p* in a human skin fibroblast line (Chen et al. 2021).

Defining novel lncRNAs, amino acid sequences for protein-coding genes, and circRNAs in HSCs provides new products that could be targets of therapy or biomarkers to assess disease progression in liver fibrosis. Products that are restricted to HSCs could provide additional specificity if targeted to inhibited fibrosis. Any unique RNA sequences contained in these products have the potential for targeting by antisense oligonucleotide approaches, and novel peptide sequences identified in protein-coding genes or produced by circRNAs have the potential to be quantified by antibody staining or could be targeted by antibodies if expressed on the cell surface. Analysis has also revealed RNA sequences that promote nuclear localization of lncRNAs in HSCs. Further understanding of the proteins that recognize these sequences for specific lncRNAs could also identify RNA binding proteins that could be modulated to disrupt the function of nuclear lncRNAs by promoting mislocalization to the cytoplasm.

In summary, we performed transcriptomic analysis to identify coding and noncoding RNA products that are at least 200 nt in length and expressed in primary human HSC myofibroblasts and used these data to define the lncRNAs and circRNAs expressed in human HSCs including transcripts that have not been annotated previously. We also expanded this analysis to include full-length transcripts allowing the identification of additional lncRNAs and un-annotated splicing events within protein-coding genes that create novel amino acid sequences. By integrating these data with datasets from patients with progressive fibrosis in NAFLD, we generated a catalog of previously-unappreciated transcripts that could help us better understand the development of liver fibrosis.

## Materials and Methods

### Cut & Tag experiments

Cut & Tag experiments were performed in duplicate using H3K4me3 antibody 13-0041 (EpiCypher) and H3K27ac antibody ab4729 (Abcam). In brief, adult HSCs were grown in DMEM (Gibco) containing 10 percent FBS. The cells were harvested using TrypLE reagent (Thermo) and were rinsed with PBS. Nuclei were extracted using NE Buffer (20mM HEPES-KOH, pH 7.9, 10mM KCl, 0.1% Triton X-100, 20% Glycerol, 0.5 mM Spermidine supplemented with 1x Roche cOmplete mini, EDTA-free protease inhibitor). The nuclei were resuspended in PBS and fixed in 0.1% formaldehyde for 2 mins. The reaction was then quenched with 1.25 M glycine. The nuclei were resuspended at 1 million nuclei per mL in wash buffer (20mM HEPES pH 7.9, 150mM NaCl, 0.5mM Spermidine supplemented with 1x Roche cOmplete mini, EDTA-free protease inhibitor) containing 10 percent DMSO, frozen in Mr. Frosty containers (Millipore) and stored in a −80°C freezer.

For each Cut & Tag reaction, 100,000 nuclei were adsorbed to activated Concanavalin A-coated magnetic beads. Nuclei-adsorbed beads were resuspended in the antibody buffer (wash buffer and 0.01% Digitonin) containing 0.5 μg antibody and allowed to incubate overnight at 4°C on a nutator. After two washing steps, the secondary antibody was added and incubated with beads for 1 hour at room temperature (RT). Beads were washed in Wash buffer 300 (Wash buffer containing 300mM NaCl) two times and resuspended in Wash buffer 300 containing Protein A/G-TN5 (EpiCypher) and incubated for an hour at RT. The tagging reaction was activated using the TN5 activation buffer (Wash buffer 300 and 10mM MgCl_2_) at 37°C for one hour. To extract tagmented fragments from nuclei, beads were dissolved in the sodium dodecyl sulfate (SDS) buffer (0.1% SDS). The tagged fragments were PCR amplified using Nextra-compatible primers to generate indexed ready-to-sequence libraries. The PCR products were cleaned up by AMPure beads (Beckman Coulter) and run on Agilent Tapestation (Agilent Technologies) for quality assessment. The cleaned PCR products were submitted for RNA sequencing.

The sequence data from Cut & Tag libraries were aligned to the human genome (hg38) using Bowtie2 (Langmead and Salzberg 2012). Duplicate reads were identified and filtered out using GATK markDuplicate script (https://gatk.broadinstitute.org/hc/en-us). The peaks were identified using SEACR package using relaxed option (Meers, Tenenbaum, and Henikoff 2019).

### Cytoplasmic and nuclei RNA fractionation

Cytoplasmic and nuclear fractions were prepared and RNA isolated as previously described (Daneshvar et al. 2020). Briefly, cells were washed two times in PBS. Cells were then resuspended in the hypotonic buffer (20 mM Tris-HCl pH 7.4, 10 mM NaCl and 3 mM MgCl_2_, 0.5% NP-40) and incubated on ice for 30 mins with a gentle vortex every 10 mins. The supernatant was collected to extract RNA representing cytoplasmic fraction using Trizol LS reagent. The nuclei pellet was resuspended in the hypotonic buffer and spun down again to rinse the nuclei. A small fraction of nuclei were examined under the microscope to confirm complete isolation. The nuclei were resuspended in Trizol reagent (Thermo Fisher) to extract nuclear RNA fraction. RNA then was extracted as outlined in the RNA extraction section. To determine whether the fractionation was successful, we performed RT-PCR to determine enrichment of *GAPDH* and *MALAT1* in nuclear RNA fraction. The RNA integrity (RIN) was then assessed using Agilent Tapestation. RNA samples with a RIN score of >9 were sequenced.

### RNA extraction

HSCs were rinsed with PBS and lysed using Trizol reagent (Thermo Fisher). The mixture was vortexed and incubated at room temperature to ensure complete lysis of cells and separation of protein-RNA complexes. RNA was then extracted using chloroform (Sigma) and precipitated using ethanol and centrifuged at 13,000 RPM for 20 min to form an RNA pellet. The RNA pellet was rinsed with 75% ethanol. The RNA was resuspended in nuclease free water and incubated with DNase I (Thermo Fisher) to remove potentially contaminating genomic DNA fragments. The RNA was then extracted using acidic phenol chloroform, precipitated in ethanol, and resuspended in nuclease free water. RNA samples with RIN score of >9 were sequenced.

### Short-read Illumnia RNA-seq

The strand-specific RNA sequencing library was prepared by using NEBNext Ultra II Directional RNA Library Prep Kit for Illumina following the manufacturer’s instructions (NEB). Briefly, the enriched RNAs were fragmented for 8 minutes at 94°C. First strand and second strand cDNA were subsequently synthesized. The second strand of cDNA was marked by incorporating dUTP during the synthesis. The cDNA fragments were adenylated at 3’-ends, and an indexed adapter was ligated to cDNA fragments. Limited cycle PCR was used for library enrichment. The incorporated dUTP in second strand cDNA quenched the amplification of second strand, which helped to preserve the strand specificity. The sequencing library was validated on the Agilent TapeStation, and quantified by using Qubit 2.0 Fluorometer (Thermo Fisher) as well as by quantitative PCR (KAPA Biosystems).

The sequencing libraries were multiplexed and clustered onto a flow cell. After clustering, the flow cell was loaded onto the Illumina HiSeq 4000 or equivalent instrument according to the manufacturer’s instructions. The samples were sequenced using 2×100bp or 2×150bp Paired End (PE) configuration. Image analysis and base calling were conducted by the HiSeq Control Software (HCS). Raw sequence data (.bcl files) generated from Illumina HiSeq was converted into fastq files and de-multiplexed using Illumina bcl2fastq 2.20 software. One mismatch was allowed for index sequence identification.

### Library preparation for long-read sequencing

Clontech SMARTer Polymerase Chain Reaction kit (Takara) was used to produce compatible polyA-transcript cDNA. The cDNA was purified using AMPure XP magnetic beads (Beckman Coulter). The SMRTbell adapters were then ligated to the cDNA product using SMRTbell Template Prep Kit (Pacific Biosciences). The library was then sequenced using PacBio Sequel II.

### Long-read data processing pipeline

Polished sequence consensus of PacBio generated BAM files was generated using the CCS program from the IsoSeq v3 pipeline (https://github.com/PacificBiosciences/IsoSeq). We merged SMRT cells from two independent runs into one movie. The Lima package was used to remove cDNA primers from the end of the reads. The consensus sequences were then refined to identify and trim polyA and concatemer sequences. The movie was then clustered using the cluster function of the isoSeq 3.1 pipeline (https://github.com/PacificBiosciences/pbbioconda). To generate higher-quality reads we then used the polish function of isoSeq workflow which in turn mapped to the human genome (hg38) using minimap2 (Li 2018). The mapped reads were processed by the TAMA suite of scripts to collapse into unique isoforms and remove potentially 5’-end degraded isoforms (Kuo et al. 2020). SQANTI3 was used on the final transcripts dataset to classify isoform structure and protein-coding potential (Tardaguila et al. 2018).

Blat program was used to identify novel protein sequences in novel isoforms discovered in HSC transcriptome (James Kent 2002). Transcripts that match the entire reference protein sequence and extend the protein sequence internally or at the terminal ends of the protein were selected for analysis.

### Transcriptome assembly and filtrations

Reads from adult and fetal HSC samples were first mapped to the human genome hg38 using STAR (ver 2.7.9a) with the following parameters --outSAMstrandField intronMotif --outSAMunmapped Within --outSAMattributes All --readFilesCommand zcat (Dobin et al. 2013). We then used StringTie (ver 2.2.0) to generate a genome-guided HSC transcriptome from mapped reads (M. Pertea et al. 2015). Transcripts that are greater than 200 nt and do not exonically overlap with protein-coding, miRNAs and translated pseudogenes are filtered using FEELnc filter argument (Wucher et al. 2017). We further filtered the transcripts based on their coding potential using CPC2 (Kang et al. 2017). GFFcompare was used to compare transcriptomes from different datasets (G. Pertea and Pertea 2020). pyGTFTK was used to process GTF files generated in manuscripts (https://github.com/dputhier/pygtftk).

### Analysis of HSC circular RNAs

To analyze and identify novel circRNAs in HSCs, we mapped the rRNA-depleted RNA-seq reads using STAR with the chimera read detection option turned on. We then filtered and annotated the resulting back splice site reads using CircExplorer2 (X.-O. Zhang et al. 2016). To identify CDS-containing circRNAs, we composed a multi-threaded python script that scans for CDSs that also takes into account the circular topology of circRNAs (https://github.com/AminMahpour/pyCircs). circIRES sequences that we use to identify potential protein expressing circRNAs were extracted using R package circRNAprofiler (Aufiero et al. 2020). We scanned the circRNA for the existence of circIRES using the Blast program (Altschul et al. 1990).

### Pseudoalignment and differential enrichment analysis

We used Kallisto to quantify transcripts in patient, cytoplasmic, and nuclear compartments. We then used the DESeq2 R package (Love, Huber, and Anders 2014) to quantify relative enrichment of each transcript in patient samples, and HSC transcripts in cytoplasmic and nuclear fractions. ggplot2 was used to create volcano plots. Heatmaps were generated with normalized transcript counts using the ComplexHeatmap package with Z-score set as the row-scaling (Gu, Eils, and Schlesner 2016).

## Acknowledgments

Portions of this research were conducted on the O2 High-Performance Compute Cluster, supported by the Research Computing Group, at Harvard Medical School. See https://it.hms.harvard.edu/our-services/research-computing for more information. ACM was supported by a Pew Biomedical Scholars Award and NIH/NIDDK R01DK116999.

---

**Figure S1.**
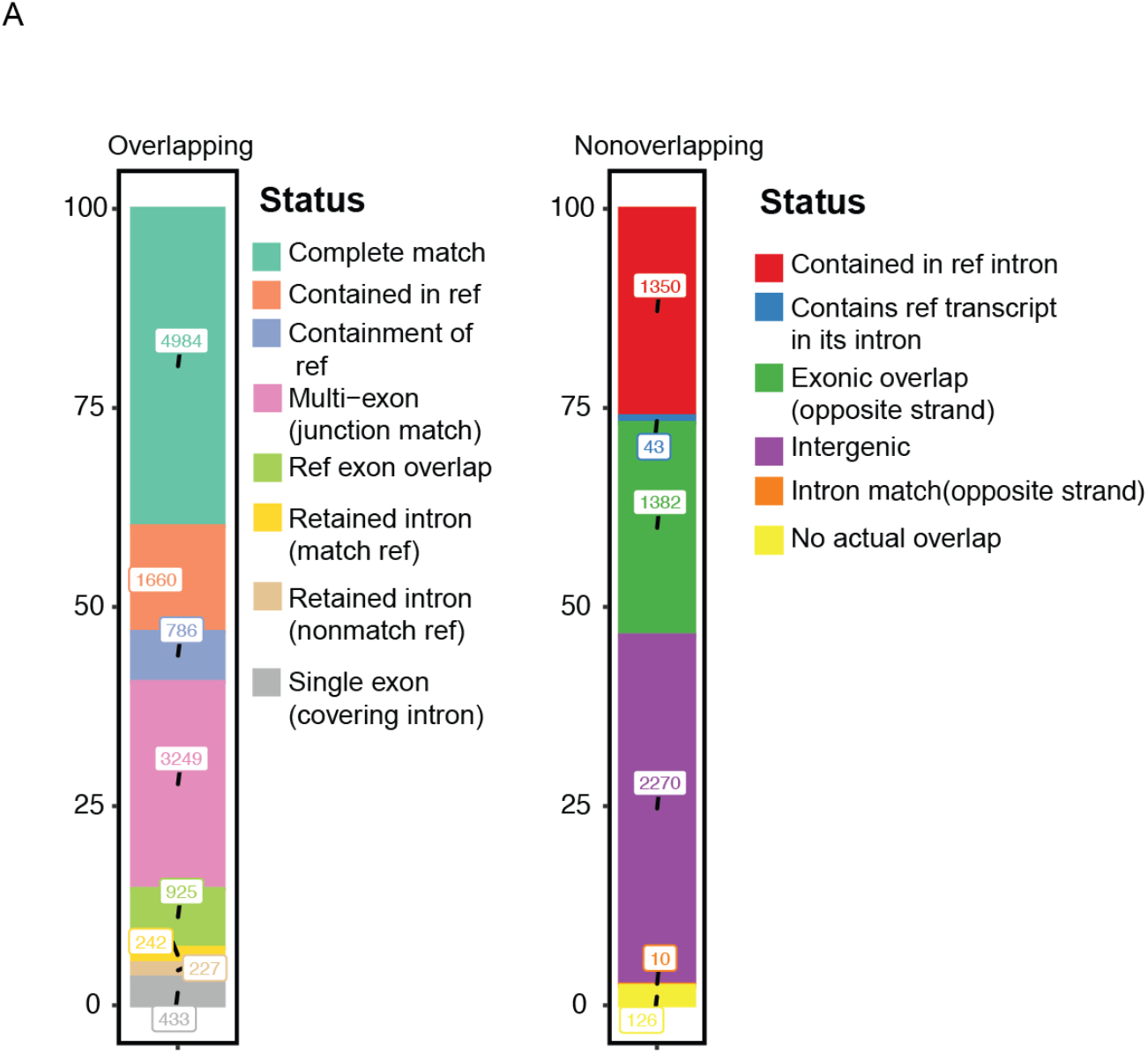
Comparison of HSC lncRNA with lncRNAs in Noncode dataset. LncRNAs that overlap with Noncode lncRNAs are shown on the left and lncRNAs that do not overlap with Noncode lncRNAs are shown on the right. The percentage in each classification is indicated on the y-axis and the number of transcripts in each category are indicated on the plots.

**Figure S2.**
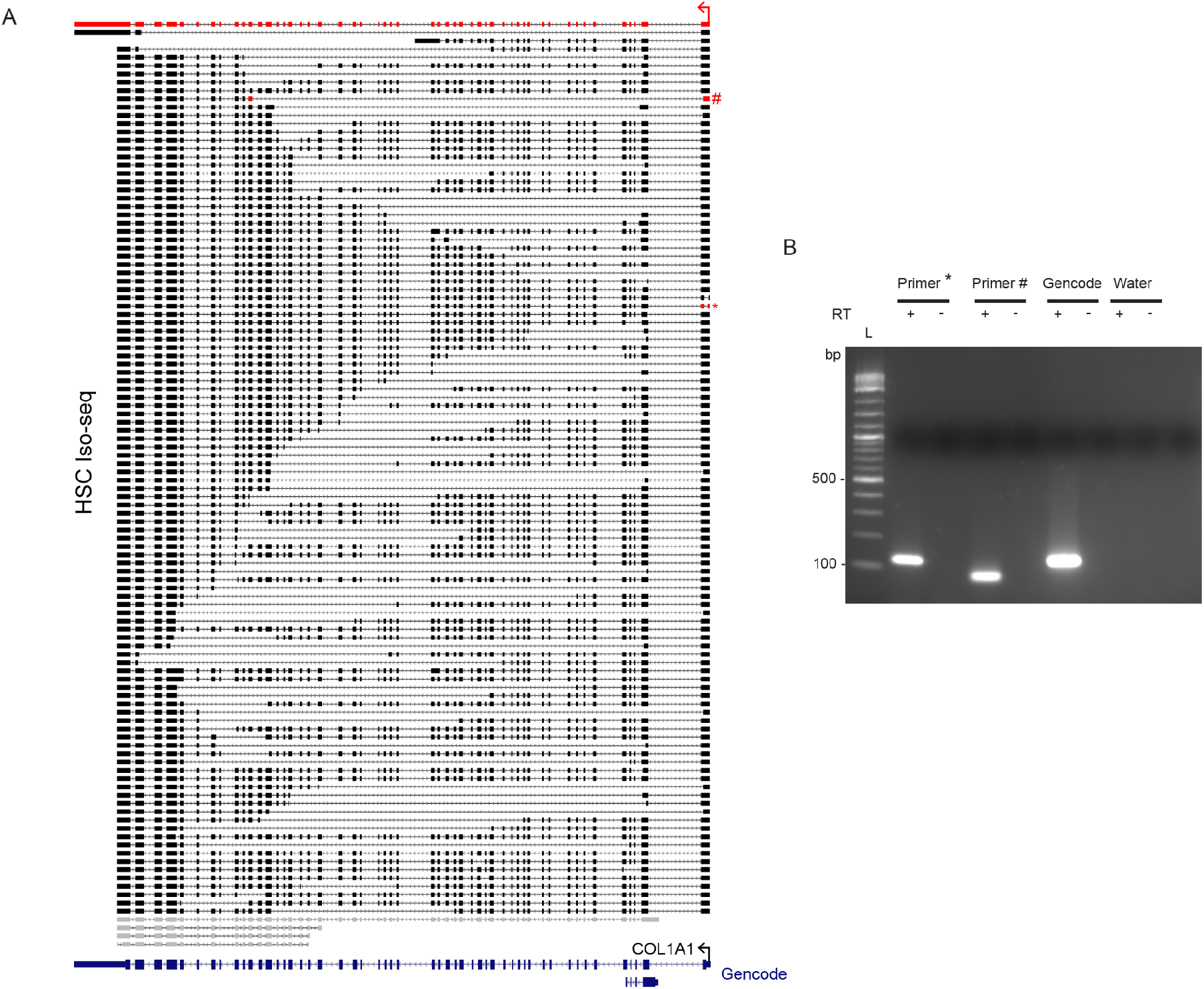
Long-read sequencing reveals transcript diversity at the *COL1A1* loci. **A)** Transcripts identified by full length sequencing analysis are displayed on the right. The annotated COL1A1 transcript is shown at the bottom and the matching isoform annotated by full-length sequencing is marked in red at the top. Gray transcripts do not share 5’-ends with the other isoforms and the reference transcript. **B)** Agarose gel image showing RT-PCR product using primers that amplify *COL1A1* isoforms in the first exon-exon boundaries as indicated by * and # red-colored symbols in (A). Three primer sets were used. Primer set * (Primer*) amplifies across the splice junction indicated by * in (A). Primer set # (Primer #) amplifies the splice junction in the transcript indicated by #. Genecode indicates primers that amplify the transcript annotated in Gencode. The plus indicates samples that underwent reverse transcription before PCR and the minus indicates samples that did not undergo reverse transcription. The lanes on the far right contain no primers as a negative control. The ladder is present in the first lane. Specific weight markers are indicated.

**Figure S3.**
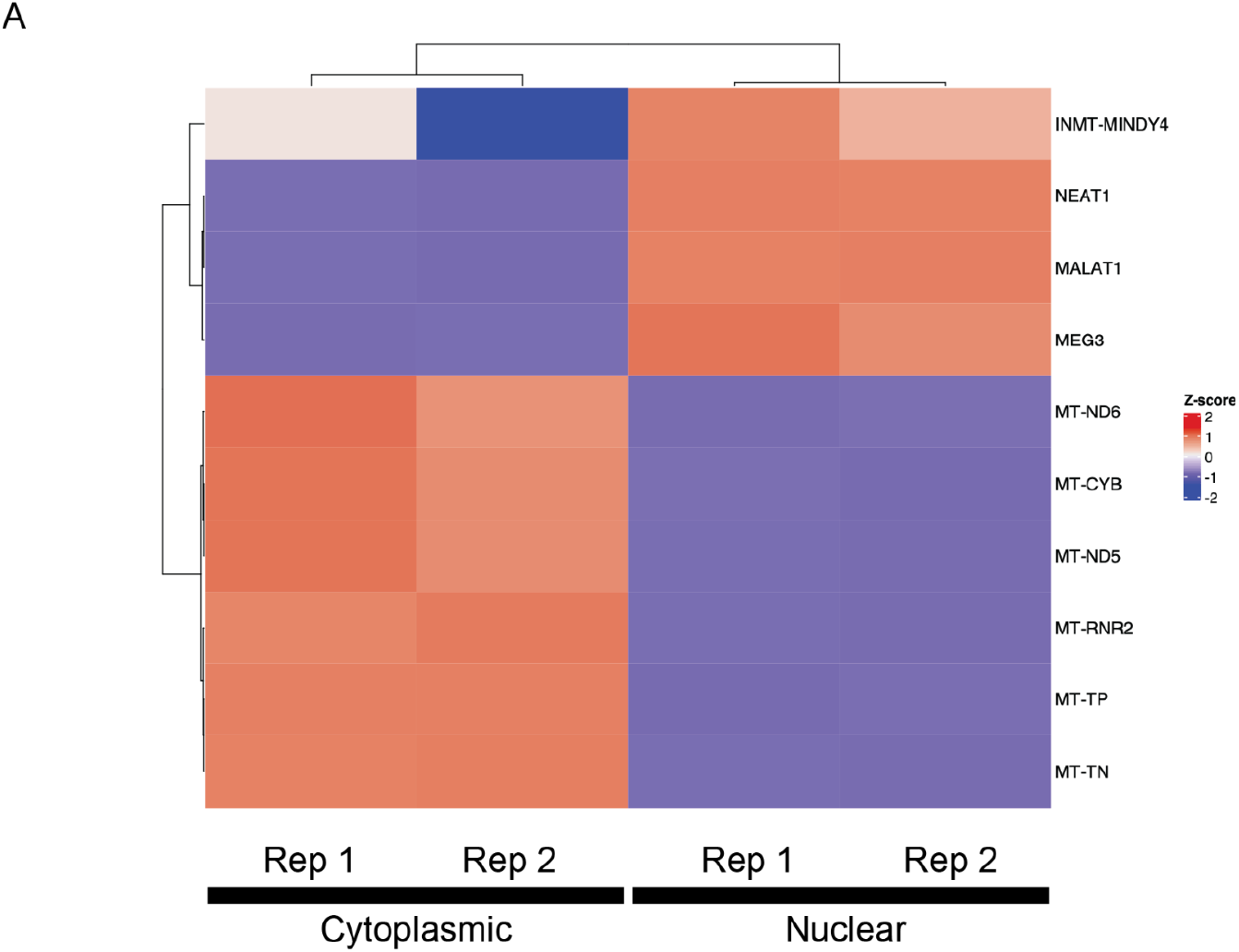
Analysis of sequencing data from nuclear and cytoplasmic fractions. Transcripts originating from the mitochondria (MT-), which should be cytoplasmic versus lncRNAs that localize to the nucleus (*NEAT1*, *MALAT1*, *MEG3*, *INMT-MINDY4*) are displayed.

**Figure S4.**
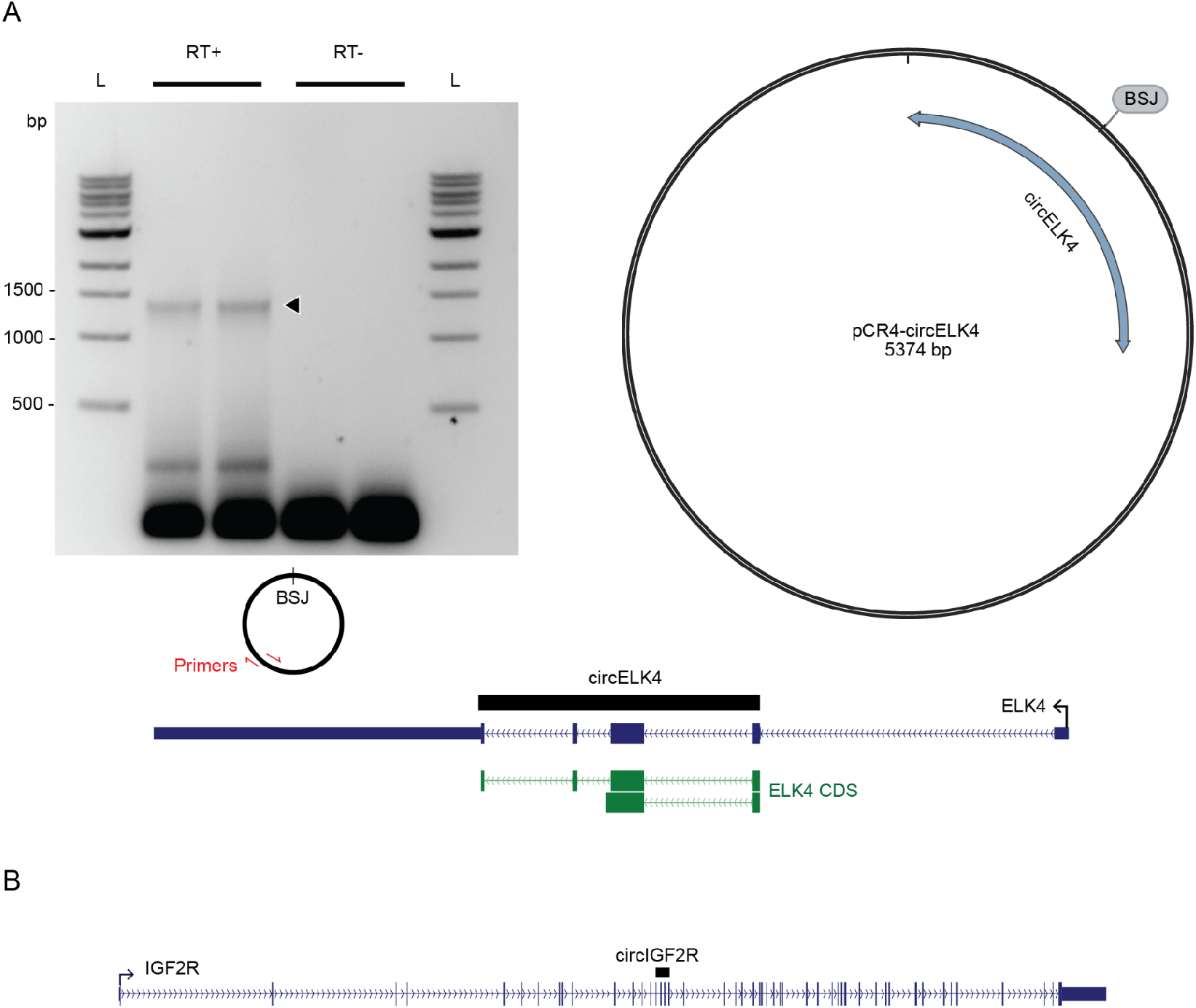
CircRNA analysis. **A)** The entire sequence of *circELK4* was amplified using divergent primers from cDNAs produced by random-hexamers as primers. The amplified sequence was then cloned into a plasmid (pCR4) for sequence confirmation. BSJ indicates the location of backsplice junction. RT+ lanes on the agarose gene underwent reverse transcription before amplification. Black arrow indicates the expected amplicon size. **B)** *circIGF2R* is produced by backsplicing of internal exons of *IGF2R* gene in HSCs and is predicted to contain 4 exons as indicated by the black rectangle above the gene.

## References

Altschul, S. F., W. Gish, W. Miller, E. W. Myers, and D. J. Lipman. 1990. “Basic Local Alignment Search Tool.” Journal of Molecular Biology 215 (3): 403–10.

Aufiero, Simona, Yolan J. Reckman, Anke J. Tijsen, Yigal M. Pinto, and Esther E. Creemers. 2020. “circRNAprofiler: An R-Based Computational Framework for the Downstream Analysis of Circular RNAs.” BMC Bioinformatics 21 (1): 164.

Banerjee, Ayan, Katherine E. Vest, Grace K. Pavlath, and Anita H. Corbett. 2017. “Nuclear Poly (A) Binding Protein 1 (PABPN1) and Matrin3 Interact in Muscle Cells and Regulate RNA Processing.” Nucleic Acids Research 45 (18): 10706–25.

Buus, Søren, Johan Rockberg, Björn Forsström, Peter Nilsson, Mathias Uhlen, and Claus Schafer-Nielsen. 2012. “High-Resolution Mapping of Linear Antibody Epitopes Using Ultrahigh-Density Peptide Microarrays.” Molecular & Cellular Proteomics: MCP 11 (12): 1790–1800.

Cabili, Moran N., Cole Trapnell, Loyal Goff, Magdalena Koziol, Barbara Tazon-Vega, Aviv Regev, and John L. Rinn. 2011. “Integrative Annotation of Human Large Intergenic Noncoding RNAs Reveals Global Properties and Specific Subclasses.” Genes & Development 25 (18): 1915–27.

Capel, B., A. Swain, S. Nicolis, A. Hacker, M. Walter, P. Koopman, P. Goodfellow, and R. Lovell-Badge. 1993. “Circular Transcripts of the Testis-Determining Gene Sry in Adult Mouse Testis.” Cell 73 (5): 1019–30.

Chen, Chun-Kan, Ran Cheng, Janos Demeter, Jin Chen, Shira Weingarten-Gabbay, Lihua Jiang, Michael P. Snyder, et al. 2021. “Structured Elements Drive Extensive Circular RNA Translation.” Molecular Cell 81 (20): 4300–4318.e13.

Creyghton, Menno P., Albert W. Cheng, G. Grant Welstead, Tristan Kooistra, Bryce W. Carey, Eveline J. Steine, Jacob Hanna, et al. 2010. “Histone H3K27ac Separates Active from Poised Enhancers and Predicts Developmental State.” Proceedings of the National Academy of Sciences of the United States of America 107 (50): 21931–36.

Daneshvar, Kaveh, M. Behfar Ardehali, Isaac A. Klein, Fu-Kai Hsieh, Arcadia J. Kratkiewicz, Amin Mahpour, Sabrina O. L. Cancelliere, et al. 2020. “lncRNA DIGIT and BRD3 Protein Form Phase-Separated Condensates to Regulate Endoderm Differentiation.” Nature Cell Biology 22 (10): 1211–22.

De Minicis, Samuele, Ekihiro Seki, Hiroshi Uchinami, Johannes Kluwe, Yonghui Zhang, David A. Brenner, and Robert F. Schwabe. 2007. “Gene Expression Profiles during Hepatic Stellate Cell Activation in Culture and in Vivo.” Gastroenterology 132 (5): 1937–46.

Derrien, Thomas, Rory Johnson, Giovanni Bussotti, Andrea Tanzer, Sarah Djebali, Hagen Tilgner, Gregory Guernec, et al. 2012. “The GENCODE v7 Catalog of Human Long Noncoding RNAs: Analysis of Their Gene Structure, Evolution, and Expression.” Genome Research 22 (9): 1775–89.

Dobin, Alexander, Carrie A. Davis, Felix Schlesinger, Jorg Drenkow, Chris Zaleski, Sonali Jha, Philippe Batut, Mark Chaisson, and Thomas R. Gingeras. 2013. “STAR: Ultrafast Universal RNA-Seq Aligner.” Bioinformatics 29 (1): 15–21.

Feng, Huijuan, Suying Bao, Mohammad Alinoor Rahman, Sebastien M. Weyn-Vanhentenryck, Aziz Khan, Justin Wong, Ankeeta Shah, Elise D. Flynn, Adrian R. Krainer, and Chaolin Zhang. 2019. “Modeling RNA-Binding Protein Specificity In Vivo by Precisely Registering Protein-RNA Crosslink Sites.” Molecular Cell 74 (6): 1189–1204.e6.

Friedman, S. L., F. J. Roll, J. Boyles, and D. M. Bissell. 1985. “Hepatic Lipocytes: The Principal Collagen-Producing Cells of Normal Rat Liver.” Proceedings of the National Academy of Sciences of the United States of America 82 (24): 8681–85.

Gao, Qiang, Yunyan Gu, Yanan Jiang, Li Fan, Zixiang Wei, Haobin Jin, Xirui Yang, et al. 2018. “Long Non-Coding RNA Gm2199 Rescues Liver Injury and Promotes Hepatocyte Proliferation through the Upregulation of ERK1/2.” Cell Death & Disease 9 (6): 602.

Guttman, Mitchell, Ido Amit, Manuel Garber, Courtney French, Michael F. Lin, David Feldser, Maite Huarte, et al. 2009. “Chromatin Signature Reveals over a Thousand Highly Conserved Large Non-Coding RNAs in Mammals.” Nature 458 (7235): 223–27.

Guttman, Mitchell, Manuel Garber, Joshua Z. Levin, Julie Donaghey, James Robinson, Xian Adiconis, Lin Fan, et al. 2010. “Ab Initio Reconstruction of Cell Type-Specific Transcriptomes in Mouse Reveals the Conserved Multi-Exonic Structure of lincRNAs.” Nature Biotechnology 28 (5): 503–10.

Guttman, Mitchell, Pamela Russell, Nicholas T. Ingolia, Jonathan S. Weissman, and Eric S. Lander. 2013. “Ribosome Profiling Provides Evidence That Large Noncoding RNAs Do Not Encode Proteins.” Cell 154 (1): 240–51.

Gu, Zuguang, Roland Eils, and Matthias Schlesner. 2016. “Complex Heatmaps Reveal Patterns and Correlations in Multidimensional Genomic Data.” Bioinformatics 32 (18): 2847–49.

James Kent, W. 2002. “BLAT—The BLAST-Like Alignment Tool.” Genome Research 12 (4): 656–64.

Jeck, William R., Jessica A. Sorrentino, Kai Wang, Michael K. Slevin, Christin E. Burd, Jinze Liu, William F. Marzluff, and Norman E. Sharpless. 2013. “Circular RNAs Are Abundant, Conserved, and Associated with ALU Repeats.” RNA 19 (2): 141–57.

Kang, Yu-Jian, De-Chang Yang, Lei Kong, Mei Hou, Yu-Qi Meng, Liping Wei, and Ge Gao. 2017. “CPC2: A Fast and Accurate Coding Potential Calculator Based on Sequence Intrinsic Features.” Nucleic Acids Research 45 (W1): W12–16.

Koch, Frederic, Romain Fenouil, Marta Gut, Pierre Cauchy, Thomas K. Albert, Joaquin Zacarias-Cabeza, Salvatore Spicuglia, et al. 2011. “Transcription Initiation Platforms and GTF Recruitment at Tissue-Specific Enhancers and Promoters.” Nature Structural & Molecular Biology 18 (8): 956–63.

Kuo, Richard I., Yuanyuan Cheng, Runxuan Zhang, John W. S. Brown, Jacqueline Smith, Alan L. Archibald, and David W. Burt. 2020. “Illuminating the Dark Side of the Human Transcriptome with Long Read Transcript Sequencing.” BMC Genomics 21 (1): 751.

Langmead, Ben, and Steven L. Salzberg. 2012. “Fast Gapped-Read Alignment with Bowtie 2.” Nature Methods 9 (4): 357–59.

Li, Heng. 2018. “Minimap2: Pairwise Alignment for Nucleotide Sequences.” Bioinformatics 34 (18): 3094–3100.

Love, Michael I., Wolfgang Huber, and Simon Anders. 2014. “Moderated Estimation of Fold Change and Dispersion for RNA-Seq Data with DESeq2.” Genome Biology 15 (12): 550.

Lubelsky, Yoav, and Igor Ulitsky. 2018. “Sequences Enriched in Alu Repeats Drive Nuclear Localization of Long RNAs in Human Cells.” Nature 555 (7694): 107–11.

Maher, J. J., and R. F. McGuire. 1990. “Extracellular Matrix Gene Expression Increases Preferentially in Rat Lipocytes and Sinusoidal Endothelial Cells during Hepatic Fibrosis in Vivo.” The Journal of Clinical Investigation 86 (5): 1641–48.

Mederacke, Ingmar, Christine C. Hsu, Juliane S. Troeger, Peter Huebener, Xueru Mu, Dianne H. Dapito, Jean-Philippe Pradere, and Robert F. Schwabe. 2013. “Fate Tracing Reveals Hepatic Stellate Cells as Dominant Contributors to Liver Fibrosis Independent of Its Aetiology.” Nature Communications 4: 2823.

Meers, Michael P., Dan Tenenbaum, and Steven Henikoff. 2019. “Peak Calling by Sparse Enrichment Analysis for CUT&RUN Chromatin Profiling.” Epigenetics & Chromatin 12 (1): 42.

Melgar, Michael F., Francis S. Collins, and Praveen Sethupathy. 2011. “Discovery of Active Enhancers through Bidirectional Expression of Short Transcripts.” Genome Biology 12 (11): R113.

Memczak, Sebastian, Marvin Jens, Antigoni Elefsinioti, Francesca Torti, Janna Krueger, Agnieszka Rybak, Luisa Maier, et al. 2013. “Circular RNAs Are a Large Class of Animal RNAs with Regulatory Potency.” Nature 495 (7441): 333–38.

Mikkelsen, Tarjei S., Manching Ku, David B. Jaffe, Biju Issac, Erez Lieberman, Georgia Giannoukos, Pablo Alvarez, et al. 2007. “Genome-Wide Maps of Chromatin State in Pluripotent and Lineage-Committed Cells.” Nature 448 (7153): 553–60.

Pantano, Lorena, George Agyapong, Yang Shen, Zhu Zhuo, Francesc Fernandez-Albert, Werner Rust, Dagmar Knebel, et al. 2021. “Molecular Characterization and Cell Type Composition Deconvolution of Fibrosis in NAFLD.” Scientific Reports 11 (1): 18045.

Pertea, G., and M. Pertea. 2020. “GFF Utilities: GffRead and GffCompare. F1000 Res. 9: 304.”

Pertea, Mihaela, Geo M. Pertea, Corina M. Antonescu, Tsung-Cheng Chang, Joshua T. Mendell, and Steven L. Salzberg. 2015. “StringTie Enables Improved Reconstruction of a Transcriptome from RNA-Seq Reads.” Nature Biotechnology 33 (3): 290–95.

Ramesh, Nandini, Sukhleen Kour, Eric N. Anderson, Dhivyaa Rajasundaram, and Udai Bhan Pandey. 2020. “RNA-Recognition Motif in Matrin-3 Mediates Neurodegeneration through Interaction with hnRNPM.” Acta Neuropathologica Communications 8 (1): 138.

Ross, Caroline Jane, Aviv Rom, Amit Spinrad, Dikla Gelbard-Solodkin, Neta Degani, and Igor Ulitsky. 2021. “Uncovering Deeply Conserved Motif Combinations in Rapidly Evolving Noncoding Sequences.” Genome Biology 22 (1): 29.

Salzman, Julia, Raymond E. Chen, Mari N. Olsen, Peter L. Wang, and Patrick O. Brown. 2013. “Cell-Type Specific Features of Circular RNA Expression.” PLoS Genetics 9 (9): e1003777.

Salzman, Julia, Charles Gawad, Peter Lincoln Wang, Norman Lacayo, and Patrick O. Brown. 2012. “Circular RNAs Are the Predominant Transcript Isoform from Hundreds of Human Genes in Diverse Cell Types.” PloS One 7 (2): e30733.

Tardaguila, Manuel, Lorena de la Fuente, Cristina Marti, Cécile Pereira, Francisco Jose Pardo-Palacios, Hector del Risco, Marc Ferrell, et al. 2018. “SQANTI: Extensive Characterization of Long-Read Transcript Sequences for Quality Control in Full-Length Transcriptome Identification and Quantification.” Genome Research. https://doi.org/10.1101/gr.222976.117.

Tørresen, Ole K., Bastiaan Star, Pablo Mier, Miguel A. Andrade-Navarro, Alex Bateman, Patryk Jarnot, Aleksandra Gruca, et al. 2019. “Tandem Repeats Lead to Sequence Assembly Errors and Impose Multi-Level Challenges for Genome and Protein Databases.” Nucleic Acids Research 47 (21): 10994–6.

Veiga, Diogo F. T., Alex Nesta, Yuqi Zhao, Anne Deslattes Mays, Richie Huynh, Robert Rossi, Te-Chia Wu, et al. 2022. “A Comprehensive Long-Read Isoform Analysis Platform and Sequencing Resource for Breast Cancer.” Science Advances 8 (3): eabg6711.

Wang, Yang, and Zefeng Wang. 2015. “Efficient Backsplicing Produces Translatable Circular mRNAs.” RNA 21 (2): 172–79.

Weirather, Jason L., Mariateresa de Cesare, Yunhao Wang, Paolo Piazza, Vittorio Sebastiano, Xiu-Jie Wang, David Buck, and Kin Fai Au. 2017. “Comprehensive Comparison of Pacific Biosciences and Oxford Nanopore Technologies and Their Applications to Transcriptome Analysis.” F1000Research 6. https://doi.org/10.12688/f1000research.10571.2.

Wucher, Valentin, Fabrice Legeai, Benoît Hédan, Guillaume Rizk, Lætitia Lagoutte, Tosso Leeb, Vidhya Jagannathan, et al. 2017. “FEELnc: A Tool for Long Non-Coding RNA Annotation and Its Application to the Dog Transcriptome.” Nucleic Acids Research 45 (8): e57.

Yang, Li, Michael O. Duff, Brenton R. Graveley, Gordon G. Carmichael, and Ling-Ling Chen. 2011. “Genomewide Characterization of Non-Polyadenylated RNAs.” Genome Biology 12 (2): R16.

Yang, Yun, Xiaojuan Fan, Miaowei Mao, Xiaowei Song, Ping Wu, Yang Zhang, Yongfeng Jin, et al. 2017. “Extensive Translation of Circular RNAs Driven by N6-Methyladenosine.” Cell Research 27 (5): 626–41.

Yu, Fujun, Jianjian Zheng, Yuqing Mao, Peihong Dong, Zhongqiu Lu, Guojun Li, Chuanyong Guo, Zhanju Liu, and Xiaoming Fan. 2015. “Long Non-Coding RNA Growth Arrest-Specific Transcript 5 (GAS5) Inhibits Liver Fibrogenesis through a Mechanism of Competing Endogenous RNA.” The Journal of Biological Chemistry 290 (47): 28286–98.

Zhang, Xiao-Ou, Rui Dong, Yang Zhang, Jia-Lin Zhang, Zheng Luo, Jun Zhang, Ling-Ling Chen, and Li Yang. 2016. “Diverse Alternative Back-Splicing and Alternative Splicing Landscape of Circular RNAs.” Genome Research 26 (9): 1277–87.

Zhang, Yang, Xiao-Ou Zhang, Tian Chen, Jian-Feng Xiang, Qing-Fei Yin, Yu-Hang Xing, Shanshan Zhu, Li Yang, and Ling-Ling Chen. 2013. “Circular Intronic Long Noncoding RNAs.” Molecular Cell 51 (6): 792–806.

Zhao, Yi, Hui Li, Shuangsang Fang, Yue Kang, Wei Wu, Yajing Hao, Ziyang Li, et al. 2016. “NONCODE 2016: An Informative and Valuable Data Source of Long Non-Coding RNAs.” Nucleic Acids Research 44 (D1): D203–8.

Zheng, Jianjian, Peihong Dong, Yuqing Mao, Shaolong Chen, Xiaoli Wu, Guojun Li, Zhongqiu Lu, and Fujun Yu. 2015. “lincRNA-p21 Inhibits Hepatic Stellate Cell Activation and Liver Fibrogenesis via p21.” FEBS Journal. https://doi.org/10.1111/febs.13544.

Zhou, Chan, Benoit Molinie, Kaveh Daneshvar, Joshua V. Pondick, Jinkai Wang, Nicholas Van Wittenberghe, Yi Xing, Cosmas C. Giallourakis, and Alan C. Mullen. 2017. “Genome-Wide Maps of m6A circRNAs Identify Widespread and Cell-Type-Specific Methylation Patterns That Are Distinct from mRNAs.” Cell Reports 20 (9): 2262–76.

